# *Where’s Whaledo*: A software toolkit for array localization of animal vocalizations

**DOI:** 10.1101/2023.08.24.554565

**Authors:** Eric R. Snyder, Alba Solsona-Berga, Simone Baumann-Pickering, Kait E. Frasier, Sean M. Wiggins, John A. Hildebrand

## Abstract

*Where’s Whaledo* is a software toolkit that uses a combination of automated processes and user interfaces to greatly accelerate the process of reconstructing animal tracks from arrays of passive acoustic recording devices. Passive acoustic localization is a non-invasive yet powerful way to contribute to species conservation. By tracking animals through their acoustic signals, important information on diving patterns, movement behavior, habitat use, and feeding dynamics can be obtained. This method is useful for helping to estimate population density, observe behavioral responses to noise, and develop potential mitigation strategies. Animal tracking using passive acoustic localization requires an acoustic array to detect signals of interest, associate detections on various receivers, and estimate the most likely source location by using the time difference of arrival (TDOA) of sounds on multiple receivers. *Where’s Whaledo* combines data from two small-aperture volumetric arrays and a variable number of individual receivers. In a case study conducted in the Tanner Basin off Southern California, we demonstrate the effectiveness of *Where’s Whaledo* in localizing groups of Cuvier’s beaked whales (*Ziphius cavirostris*). We reconstruct the tracks of six individual animals vocalizing concurrently and identify *Ziphius cavirostris* tracks despite being obscured by a large pod of vocalizing dolphins.

**Author summary:** Reconstructing the movement of animals from their vocalizations is a powerful method to observe their behavior in situations where visual monitoring is impractical. Arrays of acoustic recording devices can be used to determine the location of vocalizing animals and a series of locations can be linked to form tracks. However, reconstructing tracks requires methods of determining which animal in a group is vocalizing, finding the same vocalization on multiple recording devices, and determining the most likely location of the animal based on the relative times the sound arrived at various recording devices. We have developed a toolkit called *Where’s Whaledo* to assist researchers in reconstructing the behavior of these animals using arrays of acoustic recording devices. This toolkit greatly accelerates the process of reconstructing their tracks using a combination of automated processes and user interfaces. We use *Where’s Whaledo* to reconstruct the tracks of deep-diving Cuvier’s beaked whales. We successfully reconstruct tracks of groups of up to six whales vocalizing concurrently.

## Introduction

Passive acoustic monitoring (PAM) has increasingly been used to monitor animals in the wild [1–3]. The use of arrays of acoustic sensors has further enabled the localization of animal sounds, providing additional avenues of research including the study of behavior and a better understanding of animal population dynamics [4, 5]. Acoustic sensing has advantages over other common methods that are dependent on observers having suitable weather and lighting conditions to carry out visual surveys. PAM provides a method for non-invasive, long-term observations.

Cetaceans in particular are difficult to directly observe, but they produce species-specific vocalizations for both navigation and communication [2, 6]. Arrays of acoustic recording devices can be deployed to collect continuous data for months, providing a non-invasive method for studying cetacean behavior and presence. This method has become essential for studying deep-diving cetacean species, like beaked whales (family *Ziphiidae*), Sperm whales (*Physeter macrocephalus*), Risso’s Dolphins (*Grampus griseus*), and pilot whales (genus *Globicephala*), which are pelagic and often spend relatively little time at the surface [7–11]. PAM has provided valuable insights into their behavior despite their elusiveness [7, 12–16].

For deep-divers, PAM is emerging as an essential method for studying their population structure and dynamics [17–19]. This requires *a priori* knowledge of a number of features, like group size, vocalization rates, and acoustic detection ranges and probabilities. While some studies have estimated these parameters using acoustic models or information known about closely related species or populations, obtaining direct measurements for a specific species and site would likely improve the estimates [17]. Most of these features can be estimated by reconstructing tracks from acoustic data. Group sizes can be estimated by identifying the number of individual tracks in an encounter. Detection ranges and probabilities can be estimated based on the positions of detected animals. Additionally, passive acoustic localization can provide valuable information about depths and durations of dives, foraging depths and behaviors, responses to anthropogenic sounds or other environmental stressors, and insights into potential harm mitigation strategies.

Passive acoustic localization of cetacean vocalizations using arrays of hydrophones has been used to reconstruct tracks of a number of cetacean species, like Cuvier’s beaked whales (*Ziphius cavirostris*), common dolphins (*Delphinus delphis*), and sperm whales (*Physeter macrocephalus*)([13–16, 20–25]. Different approaches to localization have been implemented for different configurations of hydrophones, and to observe different species or behaviors of interest. Many of these studies have used localization to reconstruct two-dimensional approximations of tracks, either horizontal tracks [26, 27] or depth and range to the instrument [14, 23]. Three-dimensional localizations have been obtained using an individual hydrophone when accurate three-dimensional travel-time models could be constructed from measurements of sound speed profiles and bathymetry data [28].

Time difference of arrival (TDOA) localization uses the times a signal arrived at various receivers to estimate the location of a source. When receivers have sufficient coverage, a received signal can be localized in three-dimensional space. TDOA has been used to localize a number of vocalizing animals, including birds [29–31] bats [32], terrestrial animals [33], and aquatic animals [12, 13, 20, 34].

*Ziphius cavirostris* and *Delphinus delphis* have been tracked in three dimensions using a small-aperture volumetric array [20]. The array contained four hydrophones in a tetrahedron configuration with ∽ 0.5 m spacing between them. By measuring the TDOA between the hydrophones, the Direction Of Arrival (DOA) of the sound could be estimated as an azimuth and elevation angle to the animal. The most likely DOA was determined by minimizing the least squares error between model TDOAs and calculated TDOAs. By identifying differences in detection amplitude and azimuth angle, two individual *Z. cavirostris* whales were tracked by assuming a constant dive speed.

Localization can be performed by combining both small-aperture DOA estimates and large-aperture TDOAs [13, 35, 36]. Gassmann et al. [13] demonstrated this embedded array approach by using two small-aperture volumetric arrays and three single-channel hydrophones to localize and track *Z. cavirostris* offshore of Southern California. With these additional instruments, a total of 22 TDOAs could be used to estimate the location of a whale: six TDOAs each from two small-aperture arrays, and ten large-aperture TDOAs from five widely spaced instruments. This approach results in an overdetermined system which can improve estimation accuracy. However, uncertainty can be introduced due to ambiguous signal matching across widely spaced instruments. The difficulty increases as the number of sources increases, since the number of vocalizations arriving in the window of possible TDOAs also increases. To resolve this ambiguity, Gassmann et al. [13] plotted all possible TDOAs and manually identified the most likely correct TDOA from these sequences. They then used a maximum likelihood equation to determine the model location that best fit the measured TDOAs, successfully localizing a total of 11 individual beaked whales in groups of up to three individuals vocalizing concurrently.

Methods of associating sources automatically are necessary for accelerating the localization process. One method for source association is to temporally align sequences of clicks on widely spaced receivers [12]. If the same pattern of clicks exists in multiple hydrophones, then these patterns can be aligned to determine which clicks arrived from each source.

Automated tracking methods are emerging which use advanced multi-target tracking algorithms to identify source associations, remove false detections, and estimate likely tracks using two volumetric arrays for encounters with simultaneous detections on both arrays. [16]. Due to the directional nature of many species’ echolocation clicks [13, 37], simultaneous detections become increasingly uncommon as the distance between the instruments increases. Incorporating single-channel instruments, which are easier and cost less to deploy and recover, can increase the number of trackable encounters.

In this article, we provide a semi-automated method with opportunities for expert oversight to assist in the association of detections. We have developed a user-friendly MATLAB toolkit that builds on the methods of [20], [12], and [13] to assist researchers in obtaining tracks from acoustic datasets. To demonstrate the effectiveness of our toolkit, we used it to reconstruct ∽ 80 Cuvier’s beaked whale tracks from a four-month deployment in the Southern California Bight. We were able to reconstruct tracks for groups of up to five individuals vocalizing concurrently, a significant improvement over previous methods. We also addressed several challenges in preparing datasets for localization, including determining instrument locations and array orientations, synchronizing clocks, and calculating uncertainties. Overall, our toolkit provides an efficient tool for localizing beaked whales and other vocalizing animals and has the potential to significantly advance our understanding of their behavior and ecology.

## Methods

### Time Difference of Arrival Localization

TDOA localization is a technique that estimates the location of a single sound source by using the arrival times at which the sound is detected on multiple time-synchronized receivers. Typically the source origin time is unknown, but the difference in received times between receiver pairs can be used to determine possible source locations.

There are two forms of TDOA localization that are relevant to our process and are based on array sensor spacing: large-aperture and small-aperture. Large-aperture TDOA localization is used when the distance between the source and receivers is on the same order of magnitude as the distance between the receivers. On the other hand, small-aperture TDOA localization uses receivers that are much closer together than the distance to the source. In this case, the propagation of the signal through the arrays can be approximated as a plane wave.

#### Large Aperture TDOA

The TDOA of a signal between two receivers is determined by the distances between the source and each receiver, as shown in Eq (1).

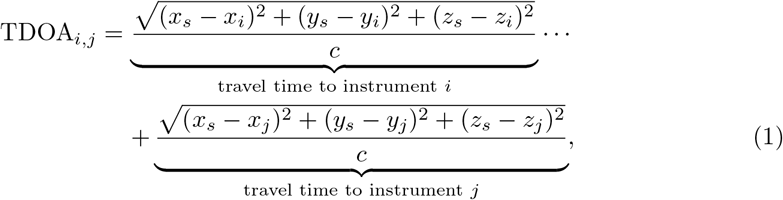

where *x*_*s*_, *y*_*s*_, and *z*_*s*_ are the Cartesian coordinates of the source location, *x*_*i*_, *y*_*i*_, *z*_*i*_, *x*_*j*_, *y*_*j*_, and *z*_*j*_ are the locations of the *i*^th^ and *j*^th^ receivers, and c is the speed of sound between the source and receivers.

The TDOA from a single pair of receivers produces a hyperboloid of potential source locations, as shown in Fig. 1A. The hyperboloid has rotational symmetry about the axis formed by the two receivers. When a detection is received on multiple receiver pairs, the source location can be estimated by finding the intersection of the hyperboloids. However, this approach works best if the receiver pairs are not collinear or somewhat orthogonal to each other and the source is interior to the region defined by the receivers.

**Fig 1.**
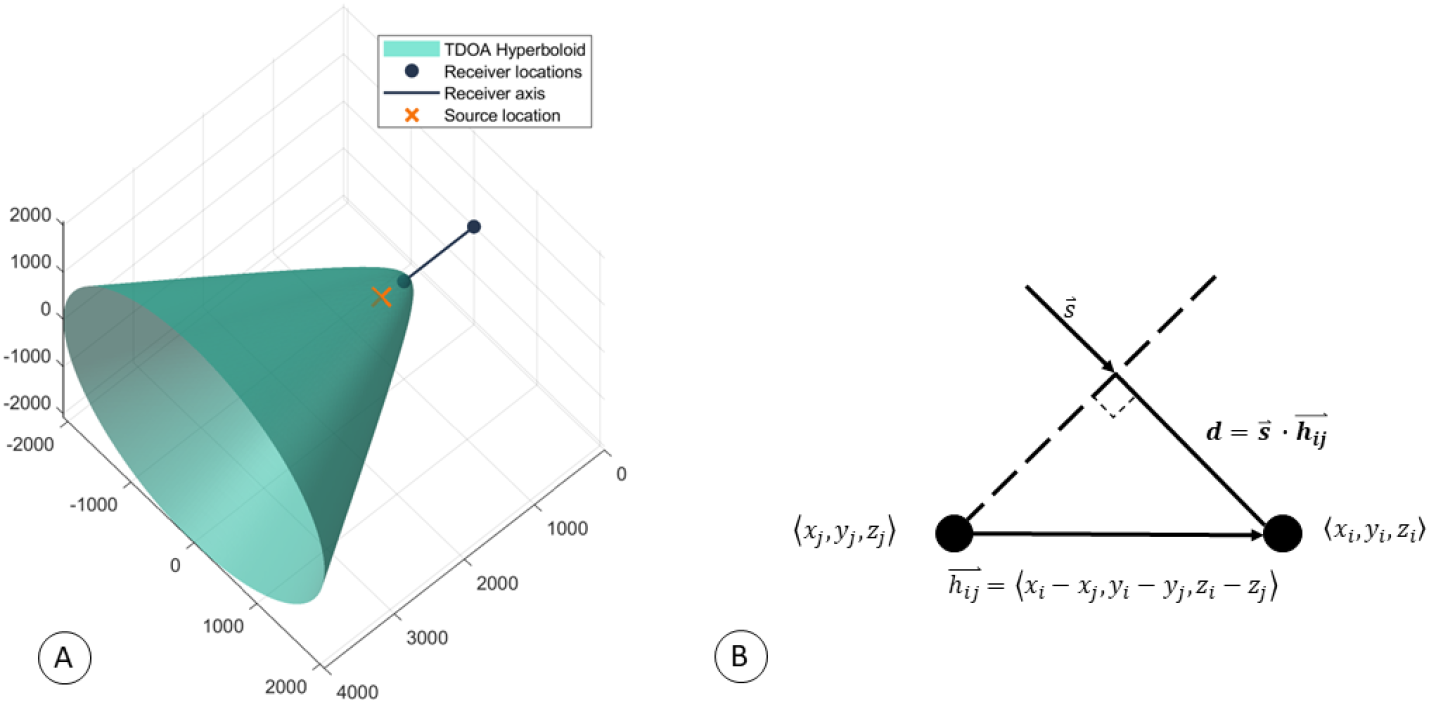
Time Difference of Arrival. Graphical representation of the TDOA for both large and small aperture separation between two sensors. A) Example of a hyperboloid of possible source locations when the TDOA between two widely spaced receivers is known. B) Small-aperture TDOA when the signal’s propagation through the array is approximated as a plane wave. The dashed line represents the wave-front, and 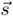 is the unit vector normal to the wavefront.

#### Small Aperture TDOA

When the distance between receivers is much smaller than the distance to the source, the calculation of the TDOA can be simplified as a plane wave propagating through the receiver array. The TDOA is the distance a plane wave travels between the receivers (d) divided by the speed of sound (c), which can be calculated as the dot product of the vector formed by the hydrophone pair 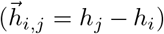 and the unit vector pointing from the source to the receiver 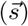. Fig. 1B and Eq. 2 below demonstrate this calculation.

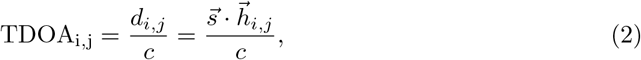

For a pair of receivers, this gives a single angle of arrival estimate, resulting in a cone of potential source locations. The hyperboloid shown in Fig. 1(A) converges to the cone formed under the plane-wave approximation and introduces negligible error. When multiple small-aperture receiver pairs are combined, the resulting cones intersect along a single line referred to as the Direction of Arrival (DOA). The DOA can be estimated from the TDOAs by placing all hydrophone pairs 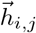 and their corresponding TDOAs into a system of linear equations, and solving for the unknown values of 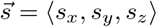 .

Since the DOA is a unit vector, it can be more intuitively represented by two angles: azimuth and elevation. We define the azimuth (az) as the top-down counter-clockwise horizontal angle, where East is 0^*°*^, and North is 90^*°*^. The elevation (el) angle is the vertical angle, where 0^*°*^ is directly down, 90^*°*^ is horizontal, and 180^*°*^ is upward toward the sea surface. We convert from 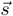 to az and el with Eqs. 3 and 4:

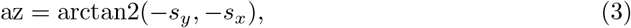

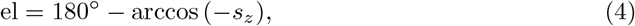

where arctan2 is the 2-argument arctangent (atan2d in MATLAB). We display these values as pointing from the receiver to the source, which accounts for the negative signs on *s*_*x*_, *s*_*y*_, and *s*_*z*_.

### *Where’s Whaledo* software package

The *Where’s Whaledo* MATLAB-based software package was designed to help analysts obtain as many animal tracks as possible by providing easy-to-use tools that allow detections to be annotated and tracks of detections to be reconstructed from localized acoustic recordings. This is done using a combination of automated processing and manual annotation of graphical data.

*Where’s Whaledo* is specifically designed to accommodate deployments with two volumetric small-aperture arrays and a variable number of single-channel receivers. To perform TDOA localization, the package provides methods to detect signals of interest, determine the time differences of a signal on various receivers, and estimate the most likely source location associated with those time differences. The *Where’s Whaledo* toolkit was built in a modular fashion, so each individual step can be adapted to obtain higher precision results or for different instrument configurations. The typical workflow is shown in Fig. 2.

**Fig 2.**
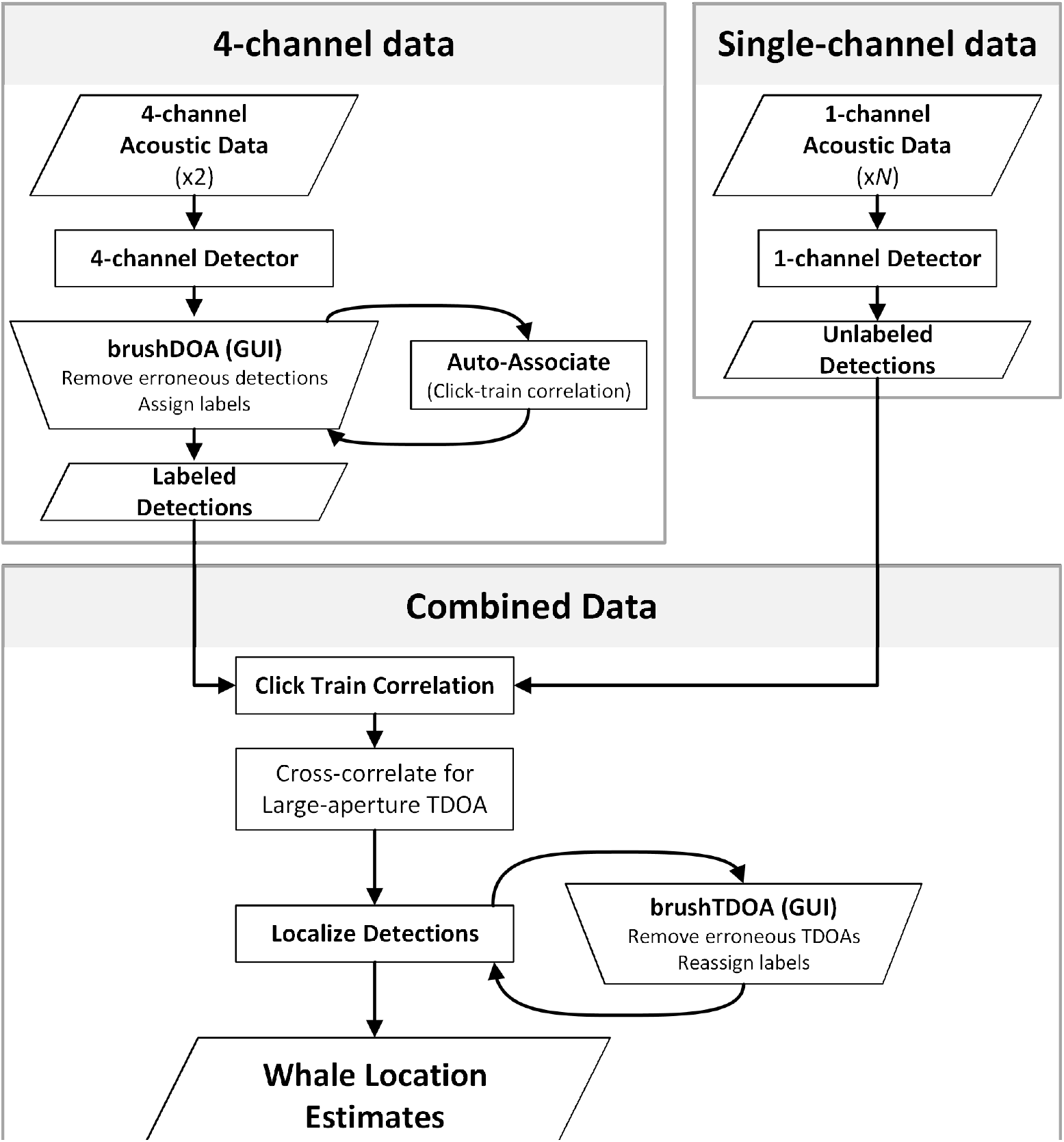
The *Where’s Whaledo* workflow. The typical workflow used to estimate whale tracks via TDOA localization. The parallelograms indicate data inputs or outputs; the rectangles represent an automated process; the trapezoids indicate a graphical user interface (GUI).

#### Detection

The sound detection step can be customized for different species based on their acoustic parameters. For detecting *Z. cavirostris* echolocation clicks, we used a fourth-order, zero-phase, high-pass elliptical filter with a cutoff frequency of 20 kHz, a peak-to-peak stop-band ripple of 0.1 dB and a minimum stop-band attenuation of 40 dB. After filtering, waveform sound pressure levels greater than approximately 68 dB re 1 *μ*Pa^2^ were identified. Peaks within a ±5 ms window around a larger peak were removed to avoid multiple cycles within a single echolocation click from being counted as separate detections. The remaining peak times were retained as potential click detections.

For the four-channel data, we cross-correlated the acoustic waveform around each detection across the other receivers in the array to determine the small-aperture TDOA. The TDOA was then converted to an azimuth and elevation using Eqs. 2, 3, and 4.

#### Association with brushDOA tool

One of the key challenges in localizing multiple sources using widely-spaced instruments is determining which detections originated from which source. To help analysts with this task, *Where’s Whaledo* used an iterative process that combines automated association with analyst manual editing using graphical representations.

A graphical user interface (GUI) tool called brushDOA was designed specifically for the purpose of removing false detections, identifying the number of unique sources, and associating detections across the two small-aperture arrays. Using this interface, an analyst can select data points to remove them from the dataset or to assign labels. Collections of detections originating from a single source can be identified by observing the gradual changes in their azimuth and elevation. If the azimuth angles from two sources are too close together to distinguish their originating sources, the elevation angles are often distant enough to provide adequate separation, and vice-versa. Analysts typically focus their efforts on labeling the array with the most detections and least ambiguity. For example, in Fig. 3, array 2 had more detections and clear separation between the various tracks, making it easier for the analyst to identify unique sources and assign labels.

**Fig 3.**
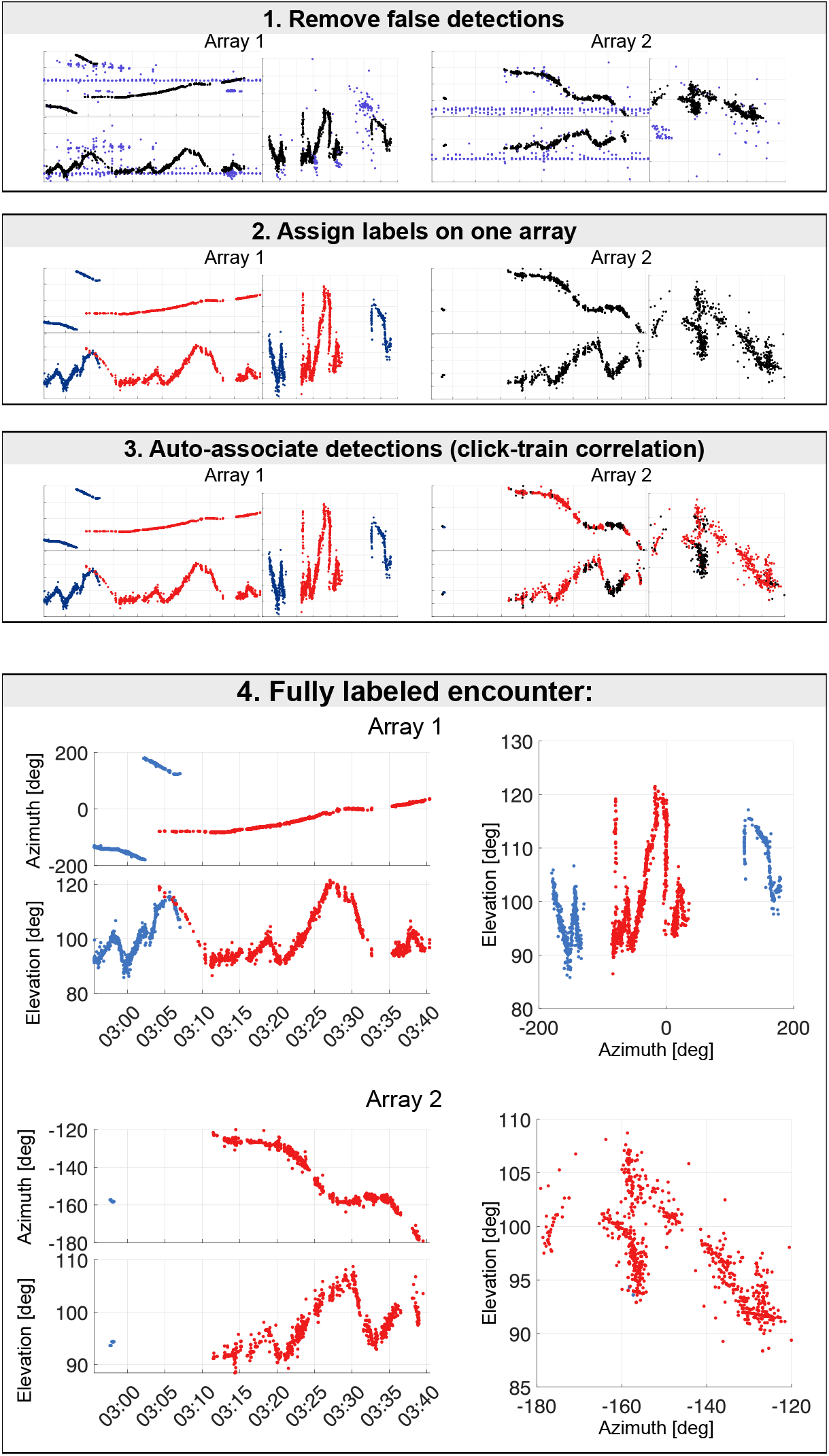
The brushDOA user interface for editing detections on two 4-channel arrays. The brushDOA user interface allows analysts to select detections, remove false detections, and assign color labels to the detections originating from the same source. The interface shows the azimuth and elevation of each detection on both arrays. The analyst removes false detections caused by other nearby sound sources (e.g. ADCP pings, dolphins, instrument noise), assigns whale labels, and runs click-train correlations to associate detections across the two arrays.

#### Association with *Click-Train Correlation* tool

After labeling one of the arrays, *Click-Train Correlation (CTC)* tool is used to associate detections across the two arrays. The CTC method identifies associations between detections from different instruments by searching for matching patterns [12]. This involves aligning a set of detected clicks in a window of time on different instruments to determine which ones originated from the same source. To accomplish this the method generates click-train vectors k_*i*_ by setting a value of one at each detection time:

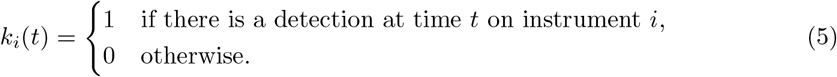

For recordings from instruments with labeled detections, different vectors of *k*_*w,i*_[*n*] are generated to include only the echolocation clicks associated with each unique label w. For unlabeled data, all detections within the window are used to create the click-train vector. Once the click-train vectors *x*_*c,i*_ are generated for each instrument and each whale, they are convolved with a 20 ms wide Hanning window to give some width to the detections. This accounts for uncertainty in the times of arrival and potential changes in the interval between clicks due to a non-stationary source. The resulting click trains are then cross-correlated, and the location of the peak of the cross-correlation between two click trains gives an estimate of the TDOA (*τ*_*w*_).

To determine which detections in the unlabeled array are associated with those in the labeled array, the unlabeled detections that align with the labeled detections after being delayed by *τ*_*w*_ are assumed to originate from the same source and are assigned labels accordingly. However, in cases where both instruments have inadequate detections from the same source, the resulting click-trains may not correlate strongly and may only produce small peaks with no clear dominant peak that can be used to estimate *τ*_*w*_. To address this, a condition is set to determine if the click-train correlation has failed due to insufficient detections arriving from the same source. Specifically, if the highest peak in the cross-correlation is not sufficiently higher than all other peaks, the click-train correlation is considered to have failed, and no detections in the window will be assigned labels. As a default parameter, if the second-highest peak is greater than 80% of the value of the highest peak, it is classified as too ambiguous. This percentage is adjustable by the user.

Once a sufficient number of detections are associated with CTC, the analyst uses them to determine which other detections are likely to have originated from the same source based on their azimuths and elevations. However, in some cases, there may be ambiguity in sources as the azimuths and elevations of two sources intersect. These sources can still be associated using CTC from the labeled detections on the other array. Once the labeling process is complete, the analyst can move on to the next phase of localization by incorporating the single-channel detections, as shown in the “Combined Data” box in Fig. 2.

The CTC function in *Where’s Whaledo* allows adjustment of several parameters including:

- the length of the window used in the click-train correlation,
- the width of the Hanning window convolved with each click train,
- the minimum ratio of the highest peak to the second highest peak in the click-train correlation required to assume the clicks are associated with the same source.

All of these parameters can be adjusted according to the instrument locations, the species of interest, and other features of a deployment. After performing click-train correlation in a window around one detection, the algorithm steps forward to the next detection and repeats the process.

Once the CTC method is used to associate animals across instruments and estimate an approximate TDOA, a fine-scale TDOA measurement is calculated by cross-correlating the acoustic data. To accomplish this, the expected detection times are used to extract the acoustic data around each detection. If there is a mismatched sampling rate, the data are resampled, then filtered and cross-correlated. The time corresponding to the peak in the cross-correlation is used as the precise large-aperture TDOA measurement.

To ensure accuracy, analysts can use a final interactive view to facilitate the removal of erroneous TDOAs or reassign labels to detections that are misassociated in previous steps. This interface is similar to brushDOA and typically requires very few changes.

#### Monte Carlo Bootstrap Localization

To improve localization accuracy, calculate confidence intervals, and combine multiple instrument pairs for each detection, a Monte Carlo Bootstrapping approach is implemented for each detection. First, small gaps in TDOAs are filled in by interpolating between recent detections. Interpolation is only performed when detections are no more than five minutes apart.

Locations are estimated using either one four-channel array and one single-channel or two four-channel arrays. For the first case, the intersection between the DOA of the four-channel and the hyperboloid formed by the large-aperture TDOA between the two instruments is found by calculating the expected large-aperture TDOA at each range step along the DOA line, then taking the range where the error between the expected and measured TDOA is minimized. When localizing with two DOAs, the source location is estimated as the point along one DOA where the distance to any point along the second DOA line is minimized.

A Monte Carlo perturbation method is used to approximate the distribution of locations that can be estimated from each set of TDOAs. Random perturbations are added to the TDOAs using a normally distributed pseudo-random number generator (randn in MATLAB) with variances of 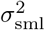 (Eq. 6) and 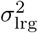 (Eq. 7) for the small- and large-aperture TDOAs respectively. The process of deriving the variances is presented in the supplemental materials. DOAs are estimated using the perturbed small-aperture TDOAs, and source locations are estimated for each combination of DOA and large-aperture TDOA available and using both DOAs. This process is repeated 50 times using different random perturbations.

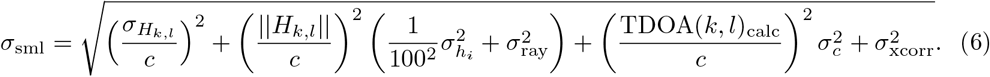

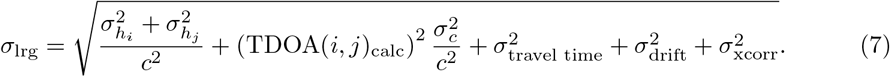

Each Monte Carlo location estimate is stored to produce a distribution of potential source locations for one detection. Location estimates are assigned a weight equal to the inverse of the variance of the location estimates using the same combination of instruments. A bootstrapping estimate of the weighted average is used to produce the final source location estimate [38, 39]. This involves randomly replacing location estimates with other estimates in the distribution and recalculating the weighted average source location estimate (resampling with replacement). Resampling is repeated 50 times, and the average of the resampled weighted average estimates is used as the final source location estimate. The 95% confidence intervals are estimated using a Studentized bootstrap method [38, 40].

#### Alternative localization approach – DOA intersect

For deployments localizing with two volumetric arrays and no single-channel instruments, the localization process can be linearized and performed much faster. The process is identical to the 4-channel data box in Fig. 2, but rather than incorporating the single-channels with click train correlation, the labeled detections from each four-channel are localized by finding the closest point of intersection between the two DOA lines. This is done by solving the system of equations relating the source location to the directions of arrival,

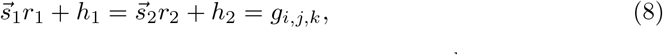

where 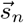 is the unit vector representing the DOA line for the *n*^th^ array, *r*_*n*_ is the range from the n^th^ array to the source, and *h*_*n*_ are the Cartesian coordinates of the *n*^th^ array location. By finding the values for *r*_1_ and *r*_2_ which minimize 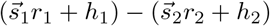, we can estimate the point along each DOA line where the lines are closest to intersecting. To do this, a 2x3 matrix 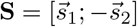 is constructed and **R** = [*r*_1_; *r*_2_] is solved for using MATLAB’s “backslash” (or mldivide) function (Eq. 9). This results in two estimates: 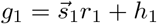 and 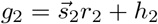 The final source estimate is the average of *g*_1_ and *g*_2_.

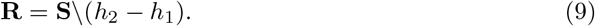

The confidence intervals for this method of localization were obtained using the jackknife variance estimator [41]. The systems of equations used to estimate *s*_1_ and *s*_2_ require only three independent TDOAs each, but these estimates are provided with three redundant TDOAs. This means one TDOA can be removed, *s*_*n*_ can be estimated with the remaining five TDOAs, and a new whale location estimate can be calculated using Eq. 9. This is repeated, removing one TDOA and localizing with the remaining 11, until 12 different whale location estimates have been produced. The variance of these location estimates is determined and used in the inverse Student’s T distribution to estimate the 95% confidence intervals.

### Case Study - Tanner Basin

We demonstrate *Where’s Whaledo* by localizing *Z. cavirostris* using a dataset collected during a four-month deployment located approximately 200 km southwest offshore of Los Angeles, California in the Tanner Basin, a region with a known presence of *Ziphius cavirostris* (Fig. 5). Four High-frequency Acoustic Recording Packages, or HARPs [20, 42] were deployed from March 16^th^ to June 11^th^, 2018. The north and south HARPs each had a single hydrophone with a sampling rate of 200 kHz moored approximately 10 m above the seafloor. The east and west HARPs each had volumetric arrays of four hydrophones in a tetrahedron configuration with ∽ 1 m spacing between hydrophones. The four-channel arrays had a sampling rate of 100 kHz and sat ∽ 6 m above the seafloor on a rigid mast. The distance between each HARP was between 470 and 1075 m.

**Fig 4.**
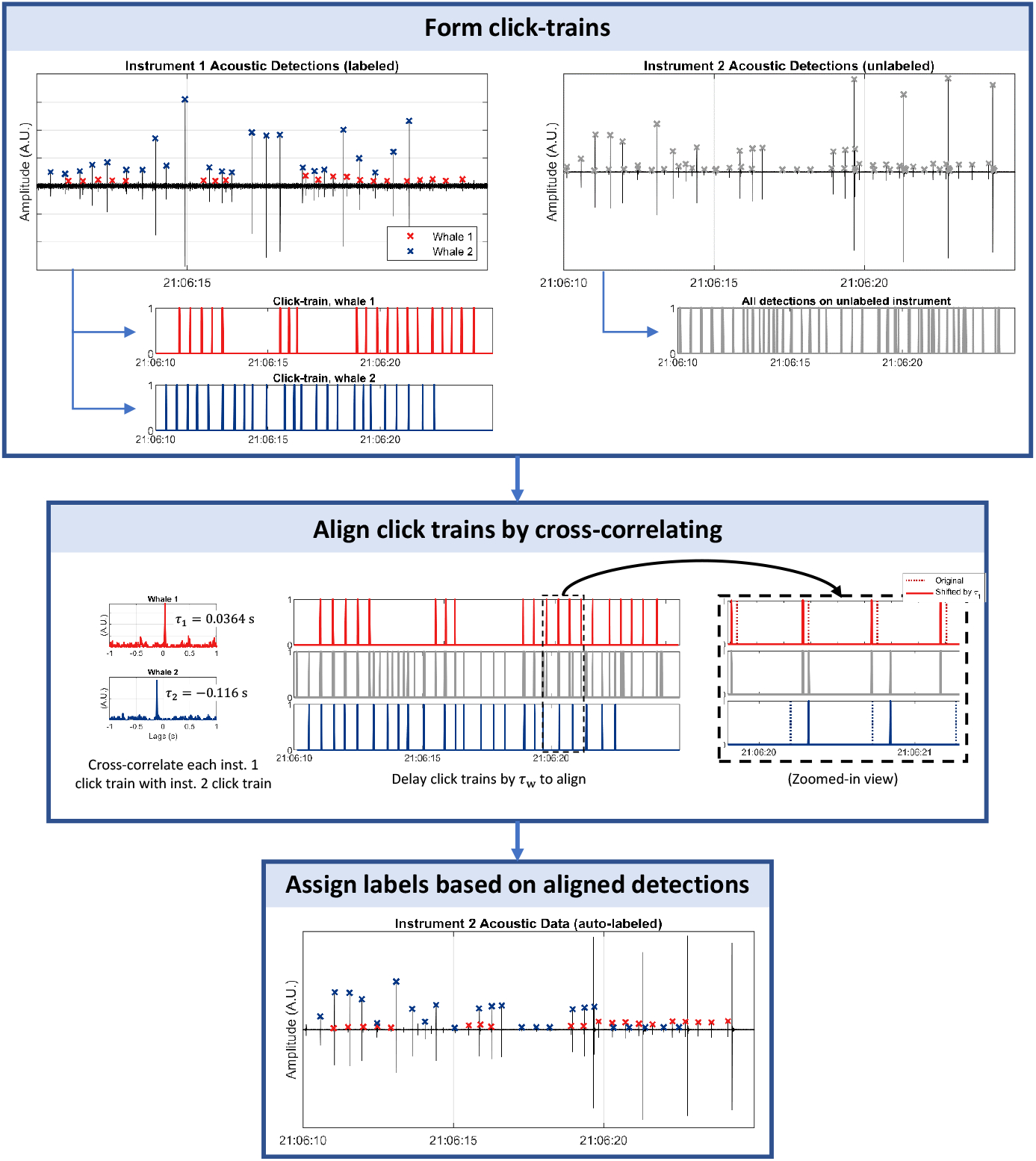
Click-Train Correlation. An example of click-train correlation (CTC) using a window of detections arriving from two sources. The labeled detections (left column) are separated into two click trains, and each is cross-correlated with the unlabeled click train. CTC is used to associate detections across instruments and determine the delay which would align the clicks.

**Fig 5.**
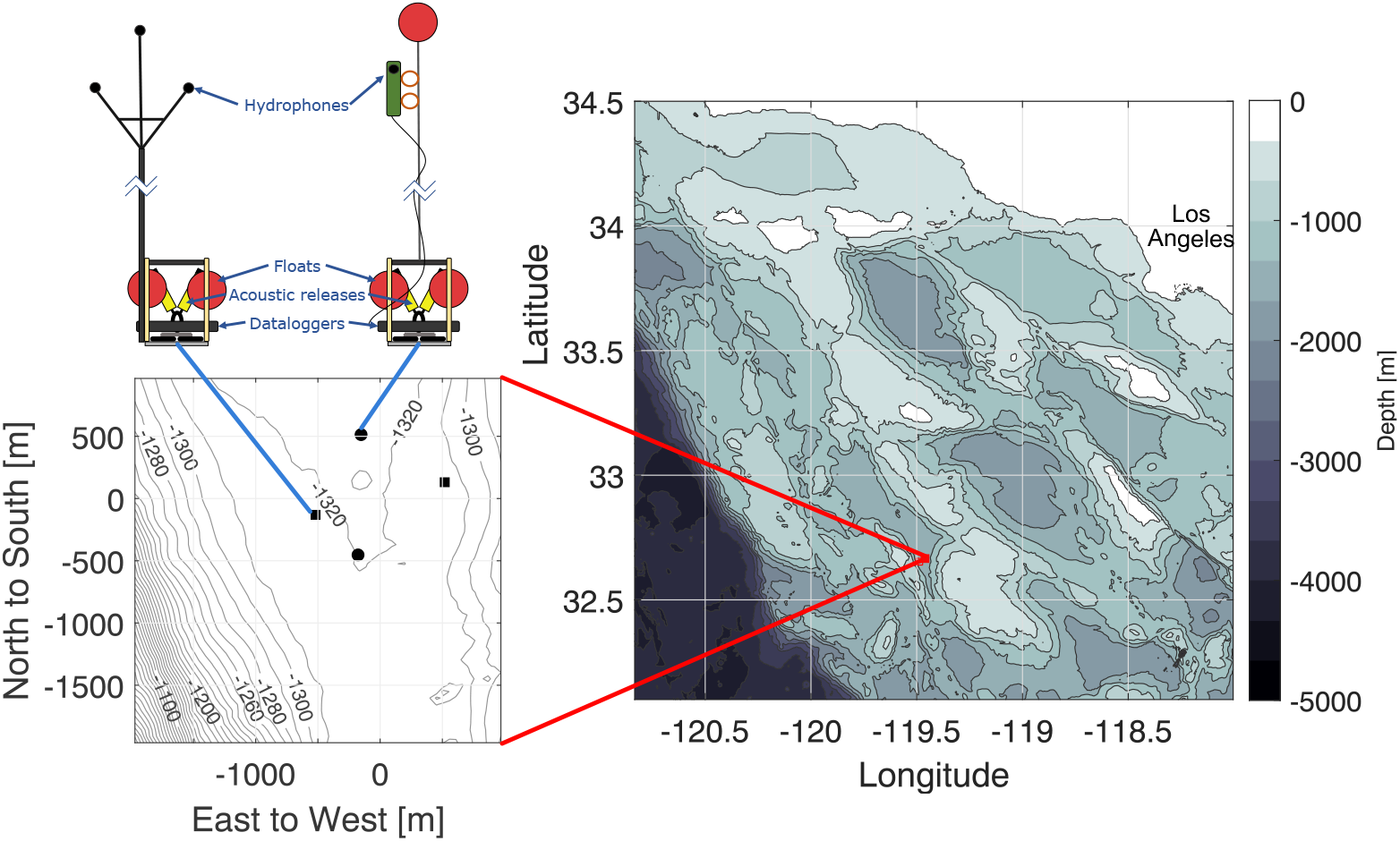
Study site. The case study site where *Z. cavirostris* tracks were reconstructed using the *Where’s Whaledo* MATLAB toolkit. Site is in Tanner Basin, ∽ 200 km southwest of Los Angeles, California. Two instrument types were used: single channel instruments (black circles on the left plot) and four-channels (black squares).

#### Oceanographic conditions and Instrument Locations

TDOA localization requires knowledge of receiver locations and the properties of the medium of propagation that affect travel times. The speed of sound in water depends on various oceanographic conditions, such as temperature, pressure, and salinity, resulting in both temporal and spatial variation in sound speed [43–45]. However, to simplify computation, a constant sound speed was used for our case study. To quantify the error introduced by this approximation, we estimated the variations in sound speed using a CTD (Conductivity, Temperature, Depth) profiler mounted near the instruments at SOCAL E. Empirical relationships between sound speed, temperature, salinity, and depth were used to estimate sound speed from the CTD measurements [43–45]. The uncertainty in the assumed sound speed is accounted for in the overall uncertainty of the localization estimates. Further details on the uncertainty calculations can be found in the supplemental materials.

Each instrument is equipped with an Edgetech acoustic release that can emit an acoustic ping in response to a ping received from a transducer on the ship. The two-way travel time of these acoustic signals from various ship locations is used to estimate the positions of the instruments. The uncertainty in instrument position is incorporated into the overall uncertainty and is discussed further in the supplemental materials.

To determine the relative positions of the hydrophones in the small-aperture arrays, we use the plane-wave approximation as shown in Eq. 2. Instead of relying on a narrow-band ping, we used the broadband engine noise emanating from the ship during the instrument localization period. The engine noise is bandpass filtered and cross-correlated to estimate the TDOA in one-second bins. The TDOA’s and the ship location for each one-second bin (obtained from the ship’s GPS system) are put into a system of equations using Eq. 2 to solve for the relative hydrophone positions within the array.

#### Clock synchronization

Ensuring clock synchronization is essential for combining data from various receivers used in localization. While all the receivers within each small aperture array were synchronized, the large aperture array required a two-step process to correct for clock drift. Initially, we synchronized the clocks using the pings transmitted by each instrument’s acoustic release during instrument localization. Then, we used the pings from an Acoustic Doppler Current Profiler (ADCP) deployed concurrently with our instruments to synchronize the clocks for the remainder of the deployment.

Each instrument’s acoustic release was enabled only during the period when it was being localized. Instrument localization was performed over the course of seven hours, and each acoustic release was enabled for between one and two hours. The pings were detected with a narrowband filter and a threshold. Due to the consistency of the amplitude of the pings, a different threshold was used for each instrument which was well above the noise levels at this frequency but had a near-zero probability of missed detection. The TDOA was calculated by cross-correlating the pings detected on each instrument. False detections produced TDOAs that significantly deviated from the true TDOAs and were manually removed. The clock drifts were calculated as the values which minimized the errors between the expected TDOAs (based on instrument locations and sound speed) and the calculated TDOAs.

The ADCP pinged approximately every minute at 75 kHz. Since the four-channel HARPs had a sampling rate of 100 kHz, the aliased frequency of 25 kHz was used to calculate the TDOA of the ADCP pings. The single-channel data were downsampled from 200 kHz to 100 kHz to deliberately alias the ADCP pings. The TDOA was then calculated by cross-correlating the detected ADCP pings for the entire deployment. The relative clock drifts between each instrument pair was then estimated as the change from the expected TDOA (based on the TDOA of the ADCP pings calculated during localization). A fifth-order polynomial fit was applied to the resulting clock drift estimates to simplify correcting for clock drift during localization.

## Results

In our case study dataset, we used a specialized beaked whale detector in tandem with DetEdit [46] to identify 600 separate time periods containing *Z. cavirostris* detections. Of these initial periods with detections, 107 contained detections with a high enough SNR and were in close enough proximity to the instruments for analysts to identify unique individuals in the encounter using `brushDOA`. However, many of these individual tracks had too few detections to be reliably localized; encounters which lasted less than 5 minutes or contained fewer than 300 detected clicks were removed from analysis, and ultimately approximately 90 encounters contained a sufficient number of localized detections in succession to be considered usable tracks. These encounters contained between one and six uniquely identifiable individual animals.

We demonstrate our approach with three examples. The first example is the simplest case, where source association is unambiguous, and tracks can be obtained quickly and easily. The second example is an encounter with six whales where source association was more challenging due to the number of whales vocalizing simultaneously and their proximity to each other. In the last example, a large pod of vocalizing dolphins obscured the beaked whale vocalizations, but we were still able to obtain tracks of two individual *Ziphius cavirostris* using DOA information and click-train correlation.

### Example 1 - Simple source association

In this example, two whales were observed that exhibited both spatial and temporal separation, facilitating a straightforward association of clicks to each source. The encounter occurred on June 11, 2018, as the whales both approached the acoustic array. The first whale swam to the northeast, passing just west of the array, while the second whale swam northward, moving directly into the center of the array (Fig. 6). The distinct spatial and temporal gap between the whales allowed for unambiguous source association. Click-train correlation performed well on both tracks, further ensuring accurate source associations. Additionally, sporadic detections of a possible third and fourth whale occurred during this time period, but they were insufficient to establish reliable track formations.

**Fig 6.**
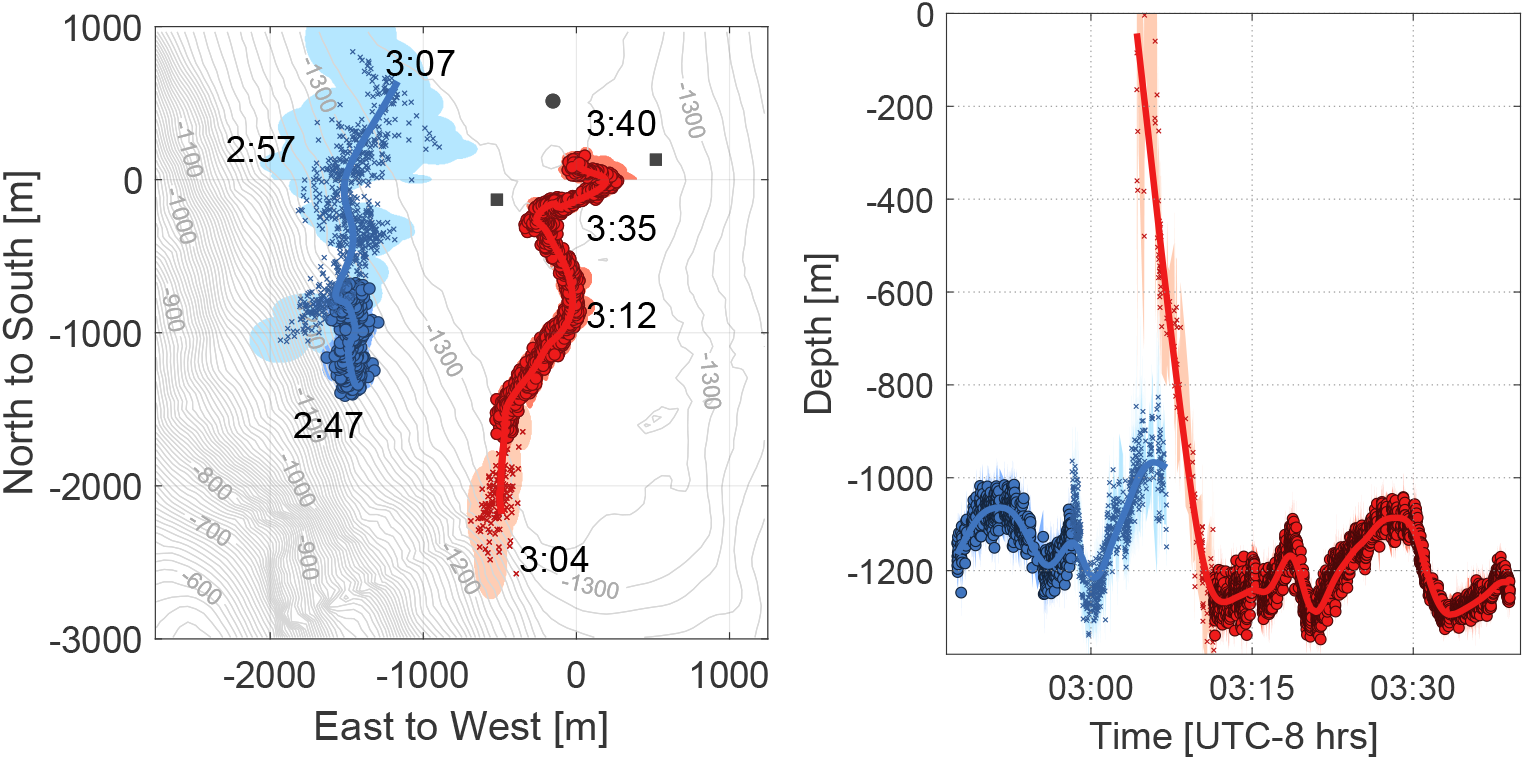
*Ziphius cavirostris* track reconstructions with clear source association. The left panel is a map view with time annotations along two separate animal tracks, and the right panel shows the animals’ depth versus time. The colors represent different whales, and the semi-transparent shading represents their 95% confidence intervals. Points with circles are localized with two four-channel instruments, whereas points with “x” were detected on only one one four-channel and one or two single-channels, Confidence intervals vary due to differences in the number of instruments used to localize, the position of the whale, or the precision and accuracy of the TDOAs.

Both whales in this example exhibited a dive descent at the beginning of their tracks. The first whale was positioned more than 3000 m from the center of the array, leading to larger depth confidence intervals compared to the second whale. During the dive phase of the second whale, detections were present only on the west 4-channel and one or both of the single channels. Once the whale reached foraging depth, all four instruments had a significant number of detections, allowing for optimal track reconstruction.

### Example 2 - Large group size

An encounter involving at least seven *Ziphius cavirostris* was detected on April 29, 2018 (Fig. 7). The whales were observed in three distinct clusters: a first group of three whales was swimming from the south and east toward the center of the array at the beginning of the encounter (red, blue, and yellow), a second pair following about 10 minutes behind from the same direction (purple and green), and a third pair swimming from the south after a 28-minute break in localizations (maroon and orange). The first two clusters are in close enough proximity in time and space that they may in fact be one group.

**Fig 7.**
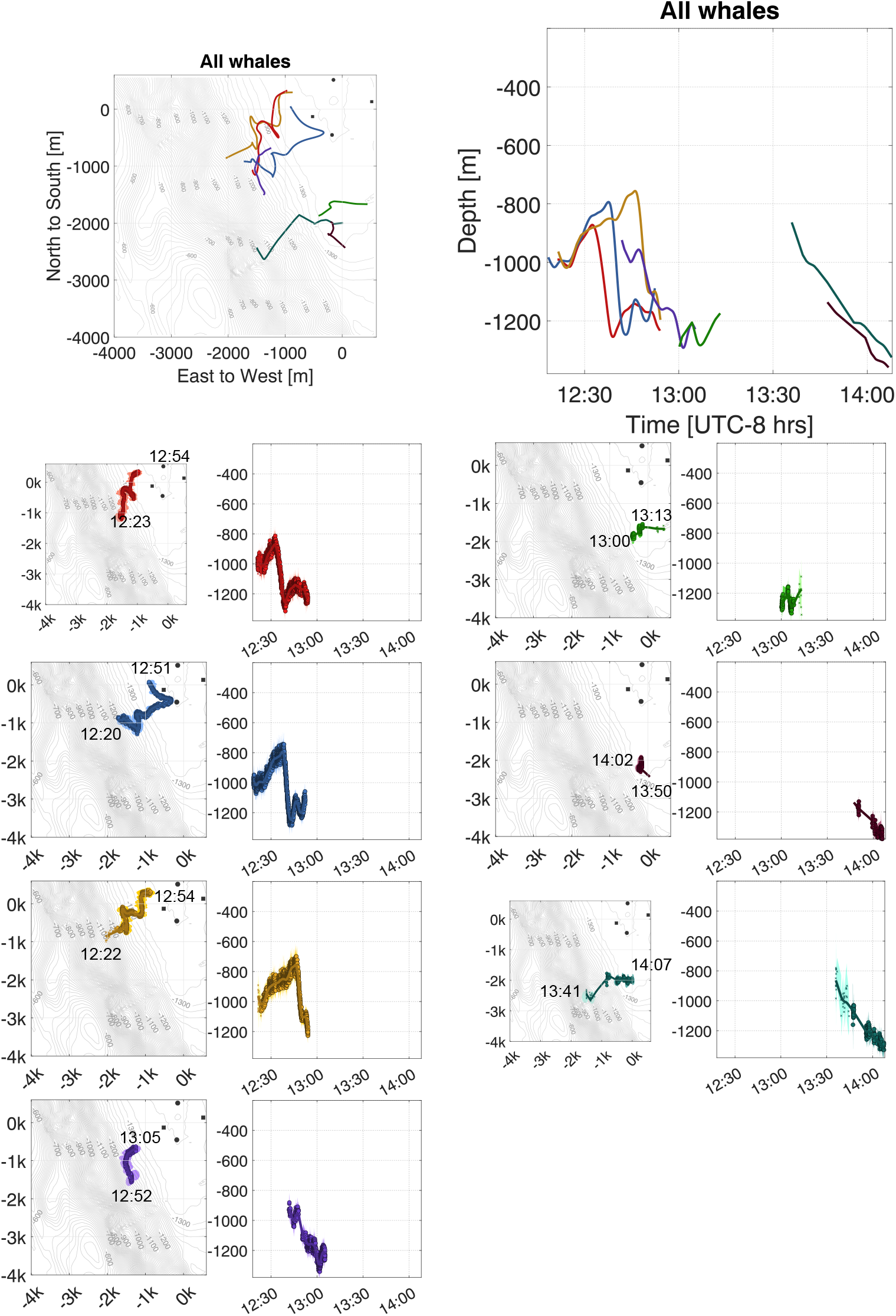
*Ziphius cavirostris* tracks with large group sizes. Top left panel shows a map view of animal tracks, and the top right panel shows the animals’ depth versus time. The lower 14 plots show the same views for each individual, where the different colors represent different whales and the semi-transparent shading represents their 95% confidence intervals.

### Example 3 - *Ziphius cavirostris* co-occurance with dolphins

An encounter on April 22, 2018 consisted of a group of two *Ziphius cavirostris* echolocating simultaneously with a large pod of dolphins (Fig. 8). It is worth noting that dolphin dive depths are much shallower than *Ziphius cavirostris*, resulting in the elevation angles of the dolphin detections being closer to 180^*°*^ than the *Ziphius cavirostris* detections. However, when the dolphin group sizes are large, as for this case, multiple individuals’ clicks arrive within the allowed small-aperture TDOAs, and their cross-correlations frequently produce erroneous DOA estimates. Consequently, the resulting DOA plots appear cluttered with detections seemingly coming from all directions, including the seafloor.

**Fig 8.**
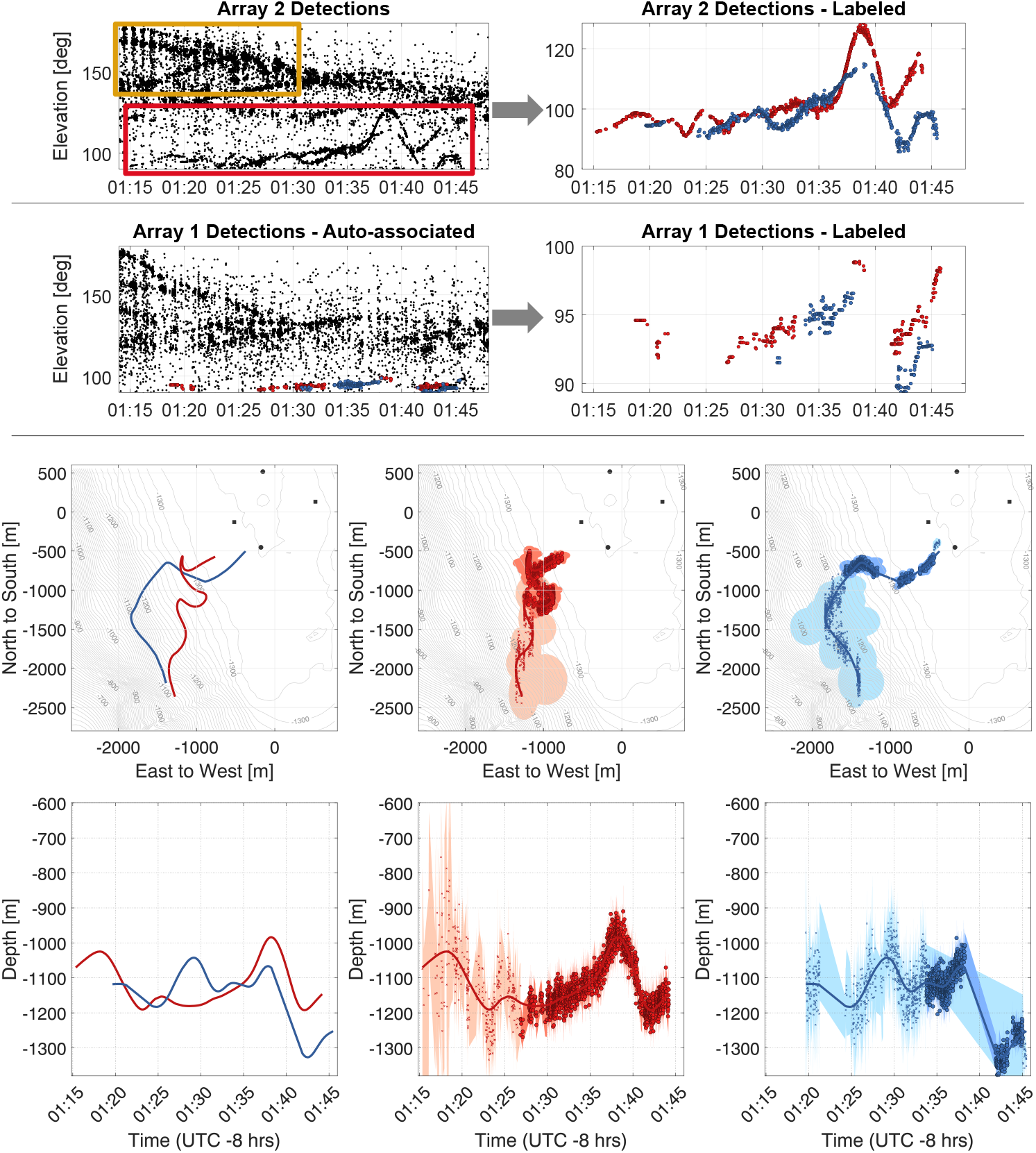
Reconstructing *Ziphius cavirostris* tracks in the presence of false-detections. Top left panel shows array-two detections, including: (yellow box) echolocating dolphins and (red box) two echolocating *Ziphius cavirostris*. Upper right panel illustrates removal of dolphin detections, due to their higher elevation angles, periodicity (where detections fade in and out on an ∽ 1 min cycle), and “fuzziness” (where multiple dolphin clicks present in one window gave erroneous DOAs). Middle panels show array-one detections (left) before and (right) after dolphin echolocation removal. Lower panels show maps with tracks of (left) both *Ziphius cavirostris* and (middle and right) individual animals.

Nevertheless, the *Ziphius cavirostris* clicks produce reliable TDOA estimates, allowing for their visual identification in the DOA plots. Dolphin detections could also be identified by their periodicity, with clicks occurring in clusters that faded in and out every few minutes. This characteristic made them easier to identify and remove from the analysis.

In this instance, identifiable *Ziphius cavirostris* tracks were present in array two, while they were less distinct in array one. Array two was therefore cleaned and labeled, followed by click-train correlation to determine the most likely *Ziphius cavirostris* clicks on array one (Fig. 8). The resulting *Ziphius cavirostris* tracks approach the array center from the southwest, apparently in a coordinated manner.

## Discussion

This study demonstrates the utility of *Where’s Whaledo* as a tool for reconstructing tracks using passive acoustic localization. We were able to obtain 90 reliable tracks from a four-month deployment offshore of Southern California. The process has the potential to be applied to similar deployments, and further development of the software could expand its usefulness to other receiver configurations, environments, and species of interest.

Identifying potential tracks and removing erroneous or unreliable detections can be done with the brushDOA GUI, which allows analysts to efficiently identify and annotate detections arriving from the same source on a small aperture array. Automated source association between widely spaced receivers is performed with click-train correlation, which searches for patterns of clicks arriving from one source in the various receivers. Once detections are correctly associated, they can be cross-correlated to determine the fine-scale TDOA, then localized using maximum likelihood comparison with a TDOA model.

Sorting clicks into those made by individual animals is the key challenge for localization. When the animals are far apart, individuals can be successfully identified in the Azimuth/elevation plots. This was occasionally challenging with *Z. cavirostris*, but for most encounters, distinct tracks could be identified on at least one of the small-aperture arrays. Calculating the TDOA on the small-aperture arrays by cross-correlating a window of time around a detection assumes only one detection within the window. For species with more individuals or whose interval between clicks is shorter than the maximum possible TDOA like some dolphin species, this may not hold, and an alternative method for identifying sources would be necessary. Click-train correlation can be effective in finding patterns of clicks on separate instruments, but may not work for other species with less unique click patterns or where detections are too sparse for adequate correlation. In these cases, analysts may rely on identifying periods of simultaneous elevation change on both arrays or incorporate other methods to associate detections with sources.

During this study, there were instances when *Z. Cavirostris* encounters coincided with the presence of delphinids, which made it challenging to track the beaked whales. One solution would be to use a more sophisticated detector that better differentiated between each species’ vocalizations, for instance using measurements of peak frequency and number of cycles within a click to separate species, or using a machine-learning based detector [47, 48]. However, due to identifiable patterns in the DOA plots, such as higher elevation angles, periodicity in vocalizations, and a high number of erroneous DOA estimates, dolphin detections were frequently able to be manually removed by analysts while *Ziphius cavirostris* detections were retained.

The tracks obtained from our approach often contain spatial offsets in clusters of detections arriving from the same source, causing the path to appear bifurcated. This is generally due to different combinations of instruments detecting the echolocation pulses;

The most reliable detections were those that were detected on both four-channel instruments. Due to the distance between the two four-channels in this deployment (1070 m) and the highly directional nature of *Ziphius cavirostris* echolocation clicks, many detections were only present on one of the four-channels. Placing the arrays closer together would increase the number of clicks detected on both four-channels. However, this would decrease the range at which reliable localizations were possible. Therefore, finding the optimal balance between the distance between the arrays and the number of clicks detected on both four-channel instruments is crucial.

To improve *Where’s Whaledo*, a more advanced detector could be used to incorporate low SNR clicks without generating false detections. Jang et al. [16] implemented a Generalized Cross-correlation detector on the same dataset, which was effective in removing most false detections caused by repeated instrument sounds. Additionally, Jang et al [16] used a multi-target tracking (MTT) algorithm to reconstruct *Ziphius cavirostris* tracks using the two small aperture volumetric arrays. Components of this algorithm could be incorporated into *Where’s Whaledo* to automate the removal of false TDOA measurements and improve source association. By incorporating estimates of an animal’s swim speed into localizations, the reliability of track reconstructions could be further enhanced [14, 16].

## Conclusion

Passive acoustic localization is a powerful way to track animal movement, which can provide valuable insights into animal behavior and the parameters needed for density and distribution measurements. A number of previous studies have demonstrated the capability of using TDOA localization of cetacean vocalizations to reconstruct their tracks. A number of challenges may limit the number of tracks obtained, including efficiently identifying potential tracks in large datasets, identifying the number of sources, and associating detections to the appropriate source. The *Where’s Whaledo* toolkit provides an efficient and reliable workflow for TDOA localization of odontocete echolocation clicks. The toolkit is designed for deployments of hydrophones containing a combination of small-aperture volumetric arrays and single-channel instruments. *Where’s Whaledo* includes a number of functions and GUIs to aid in the process of identifying separate sources, associating detections to each source, removing erroneous or unreliable detections, and estimating the most likely whale position from the TDOAs.

We demonstrate the utility of *Where’s Whaledo* by localizing *Ziphius cavirostris* echolocation clicks in the Tanner Basin. In the four-month dataset, tracks were reconstructed for ∽ 1 90 individual whales, with group sizes ranging from one to six individuals. Track reconstructions were successfully performed in the presence of significant masking due to dolphin echolocation clicks and in situations where animals were in close proximity. With some adaptations, *Where’s Whaledo* could be configured to work with a variety of receiver configurations, environments, and species of interest.

## Supporting information

**S1 Text. Derivation of TDOA uncertainties** *σ*_**sml**_ **and** *σ*_**lrg**_

## Acknowledgments

Funding provided by the Office of Naval Research Young Investigator Program, Office of Naval Research Task Force Ocean, and the Pacific Fleet. We thank Robert Headrick and Chip Johnson for their support. Special thanks to: Bruce Cornuelle for providing expertise and assistance with code implementation and matrix inversions; Junsu Jang for testing Kalman and particle filter implementations and general feedback; Bayleigh Coleman, Alma Leon, and Ryan Parkes for beta-testing earlier versions of *Where’s Whaledo* software. Ana Mae Shickich, Grace Randall, and Lauren Baggett for testing the current version of the dataset and providing valuable feedback. *Where’s Whaledo* naming credit goes to Margaret Morris.

